# TLR4-pathway impairs synaptic number and cerebrovascular functions through astrocyte activation following traumatic brain injury

**DOI:** 10.1101/2020.03.01.972158

**Authors:** Juliana M Rosa, Víctor Farre-Alins, María Cristina Ortega, Marta Navarrete, Ana Belen Lopez-Rodriguez, Alejandra Palomino-Antolin, Elena Fernandez-Lopez, Virginia Vila-del Sol, Celine Decouty, Paloma Narros-Fernandez, Diego Clemente, Javier Egea

## Abstract

**Background and purpose:** Activation of astrocytes contributes to synaptic remodeling, tissue repair and neuronal survival following traumatic brain injury (TBI). However, the mechanisms by which these cells interact to infiltrated inflammatory cells to rewire neuronal networks and repair brain functions remain poorly understood. Here, we explored how TLR4-induced astrocyte activation modified synapses and cerebrovascular integrity following TBI.

**Experimental approach:** We used pharmacological and genetic approaches to determine how functional astrocyte alterations induced by activation of TLR4-pathway in inflammatory cells regulate synapses and neurovascular unit after TBI. For that, we used calcium imaging, immunofluorescence, flow cytometry, blood-brain barrier (BBB) integrity assessment and molecular and behavioral tools.

**Key results:** Shortly after a TBI there is a recruitment of excitable and reactive astrocytes mediated by TLR4-pathway activation with detrimental effects on PSD-95/VGlut1 synaptic puncta, BBB integrity and neurological outcome. Pharmacological blockage of the TLR4-pathway with TAK242 partially reverted many of the observed effects. Synapses and BBB recovery after TAK242 administration were not observed in IP_3_R2^−/−^ mice, indicating that effects of TLR4-inhibition depend on the subsequent astrocyte activation. In addition, TBI increased the astrocytic-protein thrombospondin-1 necessary to induce a synaptic recovery in a sub-acute phase.

**Conclusions and implications:** Our data demonstrate that TLR4-mediated signaling, most probably though microglia and/or infiltrated monocyte-astrocyte communication, plays a crucial role in the TBI pathophysiology and that its inhibition prevents synaptic loss and BBB damage accelerating tissue recovery/repair, which might represent a therapeutic potential in CNS injuries and disorders.

**Declaration of transparency and scientific rigour:** This Declaration acknowledges that this paper adheres to the principles for transparent reporting and scientific rigour of preclinical research as stated in the *BJP* guidelines for Design & Analysis, Immunoblotting and Immunochemistry, and Animal Experimentation, and as recommended by funding agencies, publishers and other organisations engaged with supporting research.

**Bullet point summary:** What is already known:

- Astrocytes and microglia participate in the early cerebral and synaptic response after traumatic brain injury.
- TLR4 antagonism exerts neuroprotection in acute brain injuries.

What this study adds:

- Acute astrocyte activation contributes to synaptic loss and BBB breakdown in the acute phase of TBI, and synaptic remodeling in the sub-acute phase.
- Astrocyte activation is mediated by microglia/infiltrating-monocytes activation through TLR4 receptors.

Clinical significance:

- Inhibition of astrocyte activation through TLR4 antagonism could be a promising option for TBI treatment.

## Introduction

Traumatic brain injury (TBI) leads to a loss of function due to cell death and neuronal dysfunction in the surrounding and remote brain areas (Carron et al.2016a; Villapol et al.2014). Recent molecular to behaviour data indicate that functional recovery after TBI strongly depends on a complex network of distinct cell types including neurons, astrocytes and inflammatory cells in order to activate cell repair mechanisms, control acute inflammatory response and neurovascular integrity and initiate synaptic remodeling.

Astrocytes respond to a vast array of CNS insults and diseases (Burda et al.2016; Ceyzériat et al.2018; Clarke et al.2018) and are a major determinant of injuries’ outcome due to their role in the synaptic transmission, plasticity and neurovascular functions (Nikolakopoulou et al.2016; Perez et al.2017; Villapol et al.2014; Béchade et al.2013; Clark et al.2015; Kim et al.2016; Pascual et al.2012). Emerging evidence indicate that astrocytes exert either a neuroprotective or a neurotoxic function depending on the insult type or brain disease. In this way, neuroprotective effects of astrocytes were observed in the control of the injury size following brain injury *in vivo*; while detrimental effects on synaptic number and debris phagocytosis were reported after ischemia or infection in both *in vivo* and *in vitro* (Liddelow et al.2017; Shinozaki et al.2017). The transformation of quiescent astrocytes to a neuroprotective or neurotoxic phenotype depends on the microglia activation and release of a specific combination of cytokines and molecules of the complement component. Therefore, microglia is key to determine astrocyte fate by inducing reactive astrogliosis and by regulating astrocytic immune reactions (Donat et al.2017; Ziebell et al.2015). On the other hand, activated astrocytes-derived factors also trigger microglial activation and control their cellular functions (Jha et al.2018). In this way, microglia and astrocytes establish a bidirectional communication that is determinant for the fate of both cell types as well as for the function of neuronal networks.

Microglia and astrocytes express receptors capable of recognizing endogenous molecules released from damaged tissues and cells, which allow them to be rapidly activated upon injury. The best described of these receptors are the family of the toll-like receptors (TLRs), that serve to amplify pro-inflammatory mediator release. To date, 10 and 12 functional TLRs have been identified in humans and mice, respectively. Among them, TLR4 subtype has an important role in the progression of brain damage as mediator of blood-brain barrier (BBB) integrity and production of pro-inflammatory factors (Jiang et al., 2018; Yao et al., 2017). In the health rodent CNS, TLR4s are predominantly expressed in microglia (Bennett et al., 2016; Cahoy et al., 2008; Holm, Draeby & Owens, 2012; Zhang et al., 2014;, 2016). However, this selective expression is still debatable, as reports also show TLR4 in cultured astrocytes and neurons after injury. Nonetheless, it is clear the TLR4 expression in endothelial (Nagyoszi et al., 2010) and peripheral immune cells that enter into the brain parenchyma after damage (Frik et al., 2018).

TLR4-inhibition has been shown beneficial on the TBI progression most probably through microglia modulation of pro-inflammatory genes expression *in vivo* (Jiang et al., 2018; Yao et al., 2017). In addition, *in vitro* data show that astrocyte activation in response to TLR4 stimulation is only achievable in the presence of microglia, and that absence of functional microglia completely abolished the expression of molecular markers for astrocyte activation (Holm et al., 2012). Therefore, it is plausible that inhibition of TLR4-pathway ameliorates TBI outcome due to an indirect effect of astrocytes.

Here, we investigated how functional alterations of astrocytes induced by TLR4-pathway regulate synapses and neurovascular unit after TBI. Our data show that, in response to the damage, there is a recruitment of excitable and reactive astrocytes surrounding the injury mediated by TLR4-pathway, with detrimental effects on synapses, BBB and neurological outcome at 24-h post-injury. We also found that TLR4 activation shortly after TBI increases the release of the astrocytic-protein thrombospondin-1 (TSP-1), that serves to partially recover synapses few days after injury. Therefore, TLR4-mediated astrocyte activation plays an important role in the remodelling of neuronal networks by modulating synaptic number and controlling BBB damage, which may have important pathophysiological relevance for long-term plasticity.

## Materials and methods

### Animals

All animal experiments were carried out in accordance with the ARRIVE guidelines, the International Council for Laboratory Animal Science and the European Union 2010/63/EU Guidelines, and the recommendations made by *the British Journal of Pharmacology*. The experimental protocol was approved by the Ethical Committee for Animal Research at the Hospital Nacional de Parapléjicos (Toledo, Spain, 25-OH/2018) and by the Institutional Ethics Committee of Universidad Autónoma de Madrid, Spain (PROEX 109/18). All experiments were performed on male and female (2-3 months, 25-35-g weight) C57BL/6J wild type (Madrid, Spain. IMSR Cat# JAX:000664, RRID: IMSR_JAX:000664) and IP3R2 knockout adult mice (generously donated by Dr. J. Chen, University of California San Diego, CA, USA). IP_3_R2 knockout mice (Li et al., 2005) were backcrossed with the C57BL/6J substrain for at least 6 generations to achieve an IP_3_R2 knockout line with a C57BL/6J background. Heterozygous IP_3_R2^+/-^ mice were mate to get IP_3_R2 knockout and littermate controls. Homozygous IP_3_R2^−/−^ mice were then mate to generate a homozygous colony. Mice were genotyped by PCR analysis as described by (Li et al., 2005) from DNA extracted from tail clip showing to detect specific bands of 200 bp and 400 bp corresponding to IP_3_R2 wildtype gene and IP_3_R2 transgene, respectively. Animals were group-housed in a controlled temperature environment under a 12 h light/dark cycle, with access to food and water ad libitum. All efforts were made to minimize the number of animals used and their suffering and applying the 3Rs regulation.

### Traumatic Brain Injury Model

The Closed Head Injury (CHI) adapted from (Flierl et al., 2009) was used as a model of TBI in mice. Briefly, one hour before the procedure animals were treated with buprenorphine 0.05 mg/kg subcutaneously for analgesia. Animals were then anesthetized with isoflurane 5% and placed on a hard platform in order to minimize dissipation of energy. CHI was produced by a 50 g weight dropped from a height of 34 cm along a metal rod connected to a tip of 5 mm of diameter that hits the mice on the right hemisphere at 1 to 1.5 ML, 0 to −1 AP. After trauma, mice were immediately subjected to posttraumatic resuscitation (if necessary) and oxygen (1.5 liter O_2_/min) administration until regular breath was restored. Mice were subjected to CHI and, after post-traumatic resuscitation, animals were randomly divided into the different experimental groups. At the end of the experiment animals were sacrificed by cervical dislocation. Animals were excluded from end-point analyses if NSS was ≥ 9 one hour after brain trauma.

### Histology, Immunofluorescence and Microscopy imaging

Mice were perfused with PBS, and brains were excised and incubated in 4% paraformaldehyde for 24 hours, then in sucrose 30% until sunk before freezing in an embedding Tissue-Tek optimal cutting temperature compound (Sakura, Finetek) on dry ice. Then, 18-20 µm-thick coronal slices of frozen brains were cut with a cryostat (CM 1100; Leica), mounted on Fisher brand Superfrost/Plus microscope slides and stored at − 20 °C. Nissl staining was performed with cresyl violet (Sigma-Aldrich) as described in (Min et al., 2015). For immunofluorescence staining, slides were first incubated during 30 min at 36°C and then washed 3x in PBS, and blocked in PBS buffer containing 10% donkey serum (Jackson ImmunoResearch, West Grove, PA, USA) and 0.1% Triton X-100 (sigma, St Louis, MO, USA) for 1h. For glial fibrillary acidic protein (GFAP) and ionized calcium-binding adapter molecule-1 (Iba-1) staining, sections were incubated overnight at 4°C with the primary antibodies (Abs): 1:400 rabbit anti-GFAP (EMD Millipore catalog#ABS5804); 1:1000 rabbit anti-Iba1 (Wako catalog#019-19741). For synaptic puncta staining (post-synaptic density-95, PSD-95, and vesicular glutamate transporter-1, VGlut1), sections were incubated for two days at 4°C with the primary antibodies: 1:500 rabbit anti–PSD-95 (Invitrogen catalog#51-6900); 1:2500 guinea pig anti-VGlut1 (EMD Millipore catalog#AB230175). After incubation with primary Abs, sections were washed 3x in PBS and further incubated for 3 hours at room temperature with the respective secondary antibodies Alexa Fluor 488, Alexa Fluor 546, and Alexa Fluor 594–conjugated mouse/guinea pig and rabbit IgGs (1:500, Invitrogen). After incubation in secondary antibodies, sections were washed and incubated with 4’-6’-diamidino-2-phenylindole for 5 mins. All images of immunostained brains were collected on a Leica SP5 confocal microscope with a 63x oil immersion objective (NA = 1.35).

### Sholl analysis of microglial morphology

Z-stack confocal images from 20 μm brains sections that were immunostained for Iba-1 and DAPI were collected at 1-µm intervals using a 63x Olympus objective as described before. Images obtained from TBI ipsilateral region were collected at ∼200 µm from the damaged area to avoid necrotic damage in TBI mice. Images were obtained from three fields from each slice with two slices per mouse. Consecutive Z-stack images of the Iba-1 channel were converted to a maximum intensity projection using Fiji build of ImageJ software (RRID:SCR_002285). Images were then transformed to binary by thresholding to include microglial processes. Microglia were then isolated by manually removing surrounding processes with the eraser tool. Next, we used a line segment tool to draw a line from the center of each soma to the tip of its longest process and used the Sholl analysis plugin (Ferreira et al, 2014) to define the first shell to be around 6 μm outside of the cell body and set each step to be 2 μm size in order to determine intersections at each Sholl radius. Incomplete microglia or microglia with processes close to the margin of the field of view were avoided. From these data we determined the number of primary branches (Np, the number of branches that originated from the microglia soma). Once Sholl analysis was finished, microglia cells were manually counted in the 63x image stack to determine the mean number of microglia per field and normalized by the total number of cells that were measured using the Particle Analysis plugin in ImageJ in the DAPI channel.

### Synaptic puncta analysis

The number of colocalized synaptic puncta (VGlut1/PSD-95) was quantified as previously described (Ippolito et al., 2010). Briefly, 5-µm-thick confocal Z-stacks (optical section depth of 0.33 µm; 15 sections/Z-stack; imaged area measuring 11,272 µm^2^) of the synaptic zone in S1 cortex were imaged. Maximum projections of 3 consecutive optical sections were generated from the original Z-stack. The number of pre-, post-, and colocalized synaptic puncta was quantified by using the Puncta Analyzer plugin (developed by Barry Wark and provided by CaglaEroglu) for ImageJ software. Images were analysed by a researcher blinded to the groups. Three field of views per mouse from two sections were analysed, averaged and counted as 1. All measurements were performed in 6 independent experiments. Synaptic puncta is defined as the normalized synaptic density as the ratio of synapses in the surrounding TBI area to that in the homotopic non-injured contralateral cortex to correct for any endogenous differences in the case of transgenic mice. In the case of sham animals in which TBI was not performed, normalization was measured as the ratio of synapses in the right hemisphere to the left hemisphere.

### Calcium imaging acquisition and analysis

Animals were injected intraperitoneally (i.p.) with sulforhodamine 101 (SR101; 100 mg·kg^−1^) and left in the cage with food and water for 1–2 h (Perez-Alvarez et al., 2014). After this period, animals were anesthetized with pentobarbital (50 mg·kg^−1^, i.p.) and decapitated. Brains were quickly removed and placed in ice-cold artificial cerebrospinal fluid (ACSF) containing 124 NaCl, 2.69 KCl, 1.25 KHP_2_O_4_, 2 MgSO_4_, 26 NaHCO_3_, 2 CaCl_2_ and 10 glucose, pH 7.3 and gassed with 95% O2 / 5% CO2. Slices (350 µm) were incubated during >1 h at room temperature (21°-24° C) in ACSF. After that, the Ca^+2^ indicator Fluo-4-AM (Molecular Probes, Eugene, OR; 2–5 µl of 2 mM dye) were dropped over S1 cortex, attaining a final concentration of 2–10 µM and 0.01% of pluronic acid, for 15–20 min at room temperature (Nimmerjahn et al., 2004). In these conditions, most of the cells loaded were astrocytes, as confirmed by their electrophysiological properties in (Perez-Alvarez et al., 2014). Slices were then transferred to an immersion recording chamber and superfused with gassed ACSF in an Olympus BX50WI microscope. In some experiments TTX (1 µM) was included in the bath solution. Ca^+2^ levels in astrocytes specifically loaded with SR101 were monitored using a CCD camera attached to the microscope. Cells were illuminated with a CoolLED pE-100 fluorescent excitation system and images were acquired every 1-1.5 s using Scight software (Scientifica). Astrocytic Ca^+2^ signals were analysed offline using SARFIA, a custom-written suite of macros running in Igor Pro (Wavemetrics) (Dorostkar et al., 2010). Astrocyte Ca^+2^ levels were recorded from the astrocytes cell body and proximal ramifications and Ca^+2^ variations were estimated as changes in the fluorescence signal over the baseline. Prior to analysis, images were registered to correct movements in the *X* and *Y* directions. Movies were rejected if the plane of focus altered significantly during imaging acquisition. Regions of Interest (ROIs) containing both soma and proximal ramifications were chosen using a filtering algorithm based on a Laplace operator and segmented by applying a threshold, as described in detail in (Dorostkar et al., 2010, Rosa et al., 2015, Rosa et al., 2016). This algorithm defined most or all of the ROIs that an experienced observer would recognize by eye. Individual ROI responses were then normalized as the relative change in fluorescence (ΔF/F), smoothed by binomial Gaussian filtering, and analysed to detect activity using custom-made scripts based on a first derivative detection algorithm. A threshold set at ∼1.5-2 times the standard deviation of the time derivative trace was used to detect changes in fluorescence within the ROIs. The reliability of this algorithm to detect calcium activity was first tested by comparing the results with manual activity detection. In this paper, the fluorescent intensity of ROIs is reported as the average intensity across all pixels within its area. Fluorescent responses are reported as normalized increases as follows:

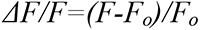

where *F* is the instantaneous fluorescence induced by a spontaneous activity and *F_o_* is the baseline fluorescence. The Ca^2+^ oscillation frequency was obtained from the number of Ca^2+^ transients occurring in 6 to 15 astrocytes in the field of view during 1 min of recording (Navarrete et al., 2012).

### Dissociation of cerebral tissue into single cell suspensions

Mice were anesthetized with pentobarbital (50 mg·kg^−1^, i.p.) and perfused with PBS. The brain was quickly removed and placed into cold Hank balanced salt solution HBSS (+Ca/Mg) medium (Gibco). The ipsilateral hemisphere was dissected out and mechanically dissected through a 100µm cell strainer. Tissue suspension was centrifuged at 286g for 5min at 4°C. Pellet was enzymatically digested in collagenase (0.16µg/mL) (Sigma-Aldrich) for 1 h at 37 °C. The cell suspension was filtered through a 70 µm filter with DNAse (666U/mL) (Sigma-Aldrich). Resuspended cell pellet in 25% of Percoll density gradient and centrifuged at 521g for 20min at 18°. The myelin coat was aspirate and cells were collected from the interface, washed once with HBSS, and processed for flow cytometry.

### Flow cytometry

After erythrocyte lysis in ACK lysis buffer (8.29 g/L NH4Cl, 1 g/ L KHCO3, 1 mM EDTA in distilled H_2_O at pH 7.4: Panreac), isolated brain cells were resuspended in staining buffer (PBS-supplemented with 10% FBS, 25 mM HEPES buffer and 2% P/S: Gibco) and their Fc receptors were blocked for 10 min at 4 °C with anti-CD16/CD32 antibodies (10 μg/mL: BD Biosciences). After blocking, cells were labeled for 30 min at 4 °C in the dark with the corresponding antibodies in staining buffer. The primary antibodies used were CD45 (Clone 30-F11, FITC, 0.6 µg/ml, Biolegend), CD11b (Clone M1/70, PercP-Cy5.5, 4 µg/ml, BD Biosciences), Ly-6G (clone AL-21, APC-Cy7, 4 µg/ml, BD Biosciences), Ly-6C (clone 1A8, BV510, 4 µg/ml, BD Biosciences), F4/80 (clone BM-8, eFluor450, 4 µg/ml, Invitrogen), CD11c (clone N418, APC, 4 µg/ml, Invitrogen) and CD284 (TLR-4, clone SA15-21, PE, 2.5 µg/ml Biolegend). After staining, cells were washed in PBS and samples were acquired on a BD FACSCanto II cytometer (BD Biosciences). Data analysis was carried out with FlowJo 10.6.2 software (Tree Star Inc., Ashaland, OR, USA). Cell type-matched fluorescence minus one (FMO) controls were used to determine the positivity of each antibody. The gating strategy employed to quantify frequencies of infiltrating and resident immune cells is shown in Fig. 7B. After cellular aggregates and debris exclusion, overall immune cells were identified based on expression of the leukocyte common antigen CD45. Different subpopulations were identified on CD45^+^ gated cells by differences in the expression levels of CD11b and CD45: Microglia was defined as CD11b^int^CD45^int^; lymphocytes were CD11b^neg^ CD45^hi^, and infiltrating-myeloid cells were defined as CD11b^hi^CD45^hi^ . The myeloid subpopulation (CD11b^hi^CD45^hi^) was further gated to quantify neutrophils (Ly-6G^hi^) and Ly-6G negative cells (Ly-6G^neg^). F4/80^+^ cells were gated on Ly-6G^neg^ subpopulation in order to distinguish macrophages. Finally, early infiltrating-monocytes were identified as Ly-6C^+^ in F4/80^+^ cells. TLR-4 expression was analyzed on resident (microglia) and infiltrating-immune cells (lymphocytes, neutrophils and infiltrating-monocytes).

**Figure 7:**
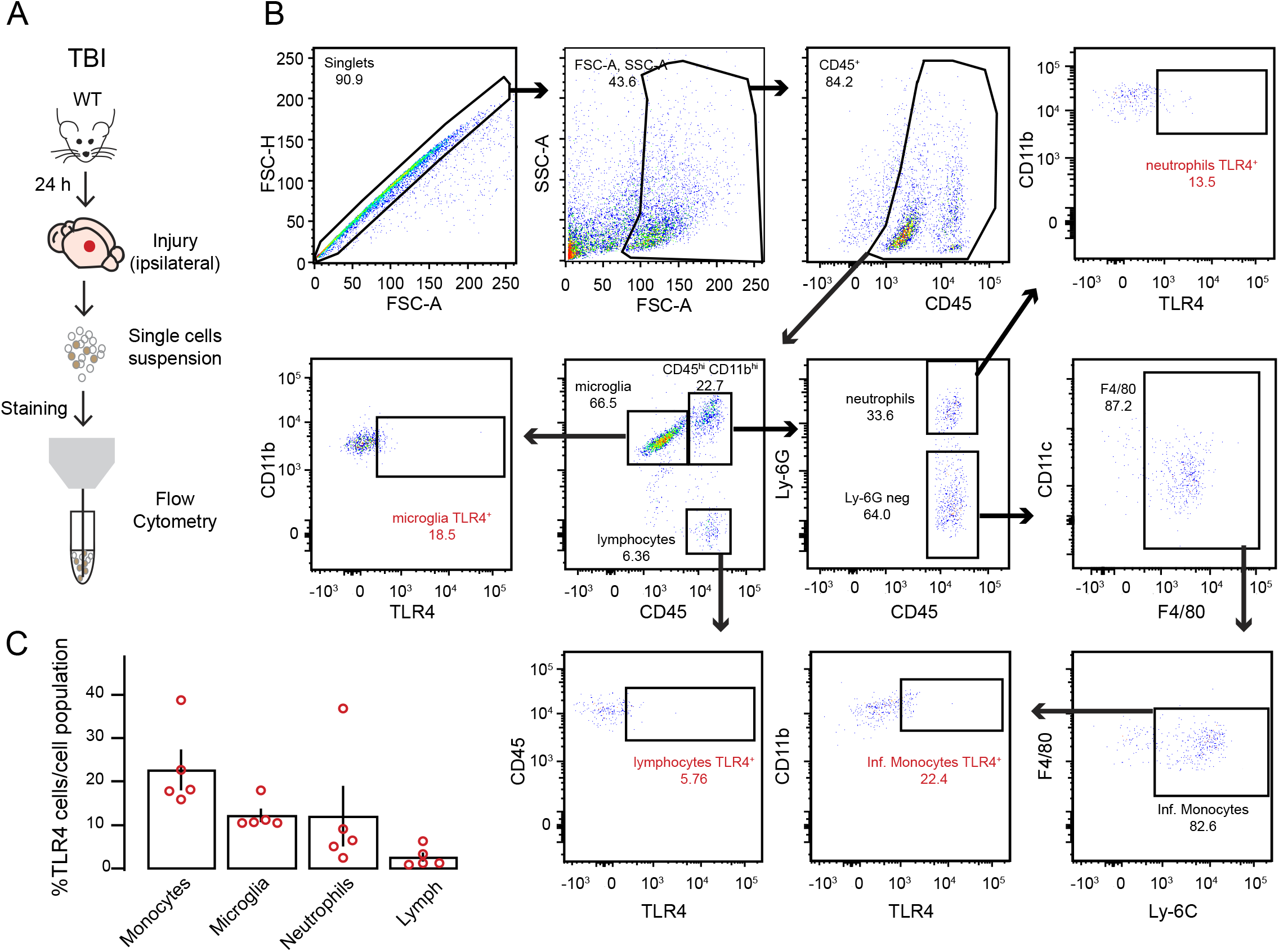
Injured TBI cortex exhibits increased infiltrating peripheral cells. (A) Schematic of the experimental protocol. (B) Flow cytometry analysis of immune cells gated on CD45^+^ from a representative TBI ipsilateral cortex. (C) Quantification of the percentage of cell types expressing TLR4 receptors of each cell population among CD45^+^ cells (microglia: CD11b^int^CD45^int^; lymphocytes: CD11b^neg^CD45^hi^; neutrophils: CD11b^hi^CD45^hi^Ly-6G^hi^ and infiltrating-monocytes: CD11b^hi^CD45^hi^Ly-6G-F4/80+Ly- 6C^hi^). Open dots on scatterplots are representative of each ipsilateral brain of n = 5 mice and depict the % of TLR4^+^ cells exhibiting extracellular receptor expression within each population.

### Neurological lesions assessment (NSS)

The severity of the injury is based on the assessment of motor and neurobehavioral functions at 1 h and 24 h after trauma using the Neurological Severity Score test (NSS), modified from (Flierl et al., 2009). The score consists of the evaluation of 10 parameters, described in Table 1. Animals with a score of 9-10 points at the 1-h test were sacrificed considering it an endpoint criterion. After the TBI, animals were kept in an isolated cage with access to food and water. One hour later, NSS was performed and points counted. Animals returned to their cages until the next day when NSS was performed once more.

**Table 1.**
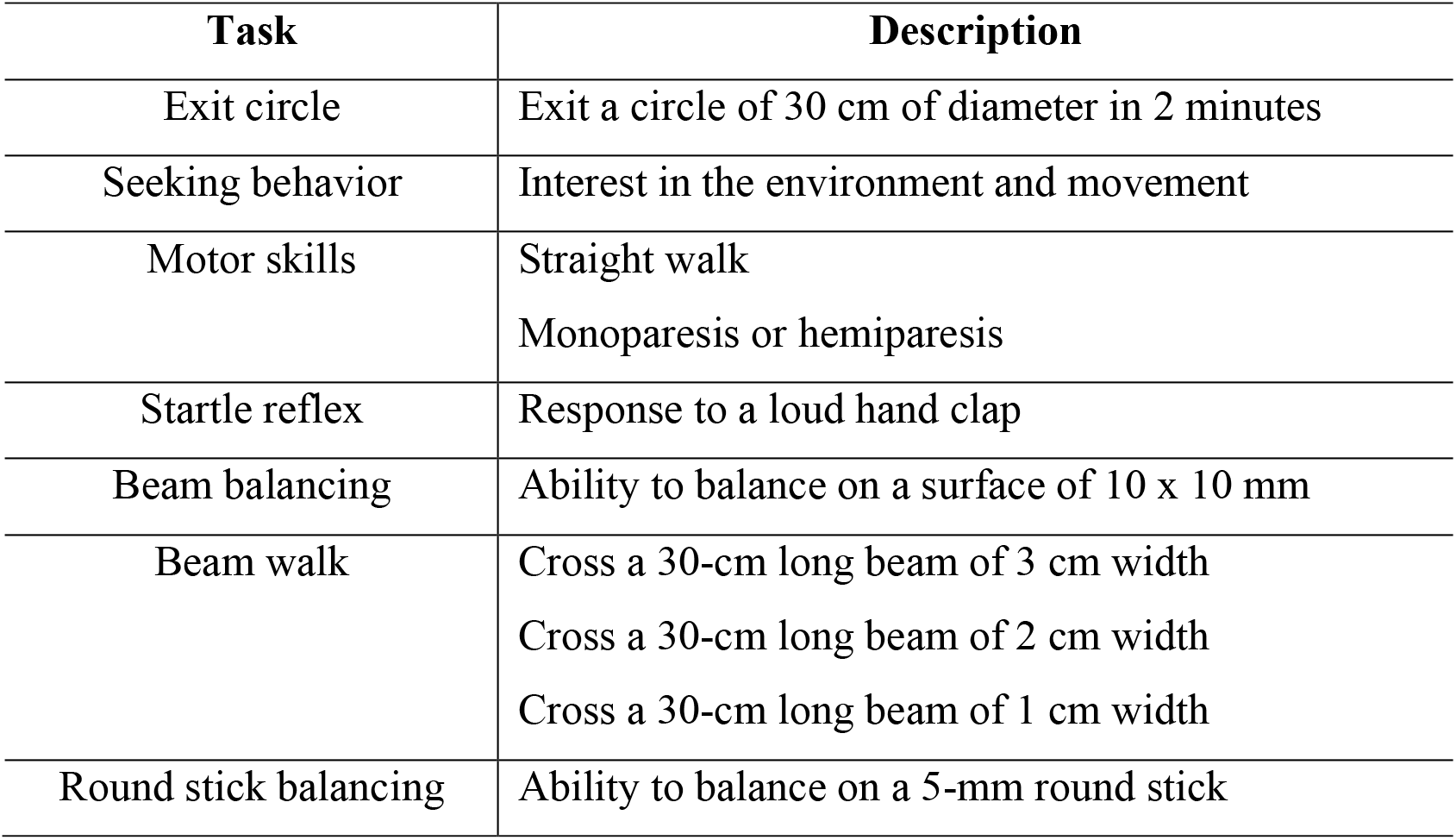
Parameters evaluated in Neurological Severity Score (NSS) test. Success in each task is considered 0 points; failure is considered 1 point.

### Pharmacological treatment

Mice were randomly divided in 3 groups: (I) sham in which mice were subjected to anesthesia but no injury was performed; (II) saline in which mice were subjected to TBI and treated with 0.9% NaCl saline solution; (III) TAK242 in which mice were subjected to TBI and treated with the TLR4 inhibitor TAK242 diluted in saline containing 3% DMSO (3 mg·kg^−1^; Sigma-Aldrich). TAK242 or saline were i.p. injected immediately following the 1-h NSS test. We were aware that TLR4 pathway is rapidly activated after a brain injury; however, in order to perform our behaviour neurological assessment without the influence of remaining levels of anesthesia, TAK242 was only administered after NSS was performed (that is, after 1 hour).

### Blood-Brain Barrier integrity

To evaluate the permeability of the blood-brain vessels and brain edema, 2% Evans Blue tracer (Sigma-Aldrich) diluted in saline was i.p. injected following the 1-hour NSS test. Animals returned to their cages and sacrificed 24h after trauma and brains were extracted and sectioned in 2-mm slices using a mouse brain slicer. A total of 4 slices were used to analyze the affected volume in the entire cerebrum. Cerebellum was not included. The slices were scanned and the total area of Evans Blue extravasation was measured in each slice using ImageJ and cubic volume (mm^3^) were calculated.

### ELISA

S100-β levels were measured in plasma using a specific ELISA kit (MyBioSource, catalog MBS2500369) following the manufacturer’s instructions. Blood was collected in the sacrifice of mice at 24 h after trauma in EDTA-treated tubes that were centrifuged at 5000 rpm during 10 minutes to obtain plasma.

### Immunoblotting

Mice brain cortex was lysed in ice-cold lysis buffer (1% Nonidet P-40, 10% glycerol, 137mM NaCl, 20mM Tris–HCl, pH 7.5, 1 g/mL leupeptin, 1 mM PMSF, 20 mMNaF, 1 mM sodium pyrophosphate, and 1 mM Na3VO4). Proteins (30 µg) from brain cell lysate were resolved by SDS–PAGE and transferred to Immobilon-P membranes (Millipore Corp.). Membranes were incubated with anti-TSP1 (1:500, NeoMarkers catalog#MS-1066-P1) or anti-βactin (1:50000, Sigma-Aldrich catalog#A3854). Appropriate peroxidase-conjugated secondary antibodies (1:10000; Santa Cruz Biotechnology) were used to detect proteins by enhanced chemiluminescence. Different band intensities corresponding to immunoblot detection of protein samples were quantified using the Scion Image program. Immunoblots correspond to a representative experiment. Immunoblots correspond to a representative experiment that was repeated four or five times with similar results.

### Data and Statistical analysis

All the data and statistical analyses comply with the recommendations on experimental design and analysis in pharmacology (Curtis et al.2018). Mice were randomly assigned to each treatment group. Data analysis of GFAP quantification, Sholl Analysis and synaptic puncta were performed by an experimenter blinded to the experimental group (that is, data were collected and named as a sequence of random numbers by one researcher, thus another researcher analysed the data completely blind to the group). All collected data were first tested with a Shapiro-Wilk test for normality and Levene’s test for equality of variance. Group measurements are expressed as mean ± standard error of the mean (SEM). To compare two groups, statistical analyses were performed using unpaired two-tailed *t*-test. Differences between more than two groups were determined by one-way ANOVA followed by Bonferroni’s *post hoc* test when there was one independent factor, or two-way ANOVA followed by Bonferroni’s *post hoc* test when there were two independent factors. The threshold for statistical significance was *p* < 0.05 throughout. Statistical differences were performed using Statistical and/or GraphPad Prism 7 (RRID:SCR_002798). Graphs and figures were made using IgorPro (Wavemetrics) and Adobe Illustrator. All measurements were undertaken in at least three technical replicates.

### Nomenclature of targets and ligands

Key protein targets and ligands in this article are hyperlinked to corresponding entries in http://www.guidetopharmacology.org, the common portal for data from the IUPHAR/BPS Guide to PHARMACOLOGY (Harding et al.2018), and are permanently archived in the Concise Guide to PHARMACOLOGY 2015/16 (Alexander et al.2019).

## Results

### TLR4 antagonism reduces astrogliosis and microgliosis after TBI

We first determined how TLR4-pathway regulates astrocyte reactivity and induction of pro-inflammatory microglia in the CHI mice model of TBI that simulates a biomechanics similar to human TBI (Albert-Weißenberger et al.2012; Flierl et al.2009). For that, we pharmacologically blocked TLR4-receptors by treating TBI animals with the selective TLR4-antagonist TAK242 (3 mg·kg^−1^, Fig. 1A) or vehicle 1 hour after injury. Cresyl violet-stained showed substantial cortical damage at 1 day after injury (1 d.a.i., Fig. 1A) following a moderate unilateral injury over the somatosensory (S1) and motor cortical areas. The cortical damage was composed of an injury core (primary damage, absence of Nissl bodies, Fig. 1a’’ region 1) surrounded by a peri-impact area (secondary damage, loss of several Nissl bodies, Fig. 1a’’ region 2) with no overt subcortical damage. TBI and sham brains were then immunostained against GFAP or Iba-1 to mark reactive astrocyte or microglia, respectively (Fig. 1B-D), and quantification taken from 200-400 µm from the injury core (Fig. 1a’’ region 3) to avoid the area with ongoing neuronal death. Increase number of GFAP^+^ astrocytes was observed in the ipsilateral cortex, while only few GFAP^+^ cells were found in sham or contralateral cortex (Fig. 1C), corroborating previous reports showing absence of astrogliosis in cortical areas with no apparent damage (Shinozaki et al.2017). TAK242 treatment significantly reduced GFAP^+^ cells (one-way ANOVA F_(4,33)_ = 81.45, p < 0.001). To examine by which degree microglia enter in a pro-inflammatory state after CHI (Wu et al.2014), we analysed their morphology in two ways. First, we counted the number of microglia displaying pro-inflammatory phenotype, that is, large cell bodies and shorter and thicker projections (hypertrophic type) or enlarged cell bodies with multiple short processes forming thick bundles (bushy type), or non-inflammatory phenotype, that is, ramified with small cell bodies and thin projections (Fig. 1B). The decreased number of the ramified phenotype with consequent increase in the pro-inflammatory one observed in the TBI ipsilateral cortex was significantly affected by TAK242 treatment (Fig. 1D, non-inflammatory: one-way ANOVA F_(4,22)_ = 4.44, p =0.008; pro-inflammatory: one-way ANOVA F_(4,22)_ = 24.76, p < 0.001). We then performed a Sholl Analysis to provide a more accurate measurement of microglia activation. Figure 1E summarizing the Sholl analysis plots of distinct groups shows that the branching profile of microglia in the region nearest the damaged tissue is shifted to the left, indicating a decrease in branching as a function of distance from the soma. Figure 1F shows the effect of TAK treatment on the number of microglia primary branches calculated by the Sholl analysis. The results of two-way ANOVA revealed a significant effect of hemisphere (F_(1,22)_ = 4.60; p = 0.043), treatment (F_(1,22)_ = 21.85; p < 0.001), and interaction (F_(1,22)_ = 5.52; p = 0.028), indicating that TAK treatment exhibited a significant effect on microglia activation.

**Figure 1:**
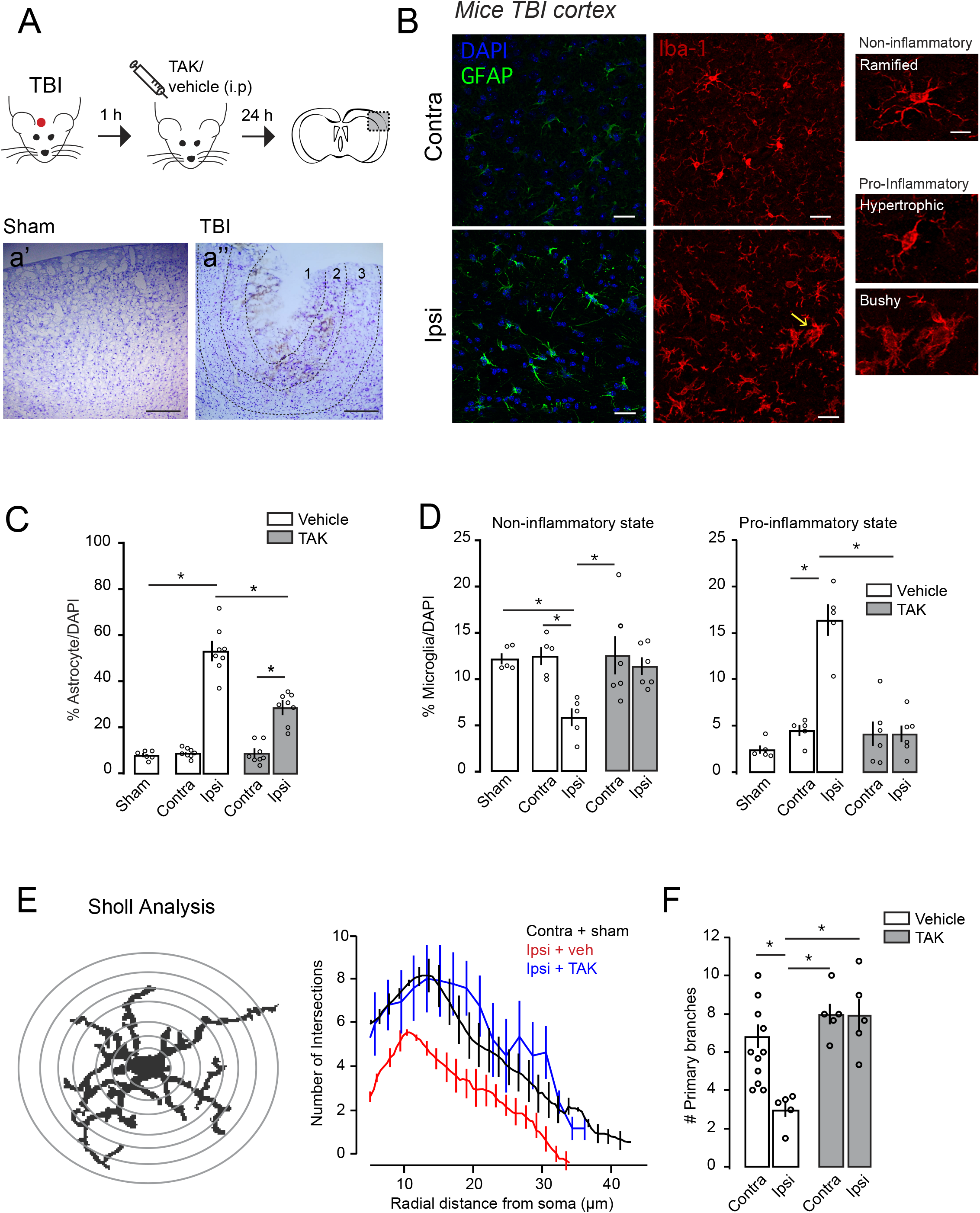
TLR4 antagonism reduces astrogliosis and microgliosis caused by traumatic brain injury. (A) Top: Schematic diagram showing the experimental strategy for the injury, pharmacological treatment and brain slice (adapted from (Paxinos & Franklin2019)). TAK242 or vehicle were i.p. injected 1 hour after injury. Dotted black box represents the area in the somatosensory (S1) cortex where functional and anatomical experiments are performed. Images in a’ and a’’ are from the area delimited in the dotted black rectangle. Nissl staining tissues showing gross morphological damage 24 h after sham (a’) or TBI (a’’) injury. Scale bar: 150 µm. Dotted lines in a’’ delimits of Region 1 (∼200 µm), which represents the injury core, Region 2 (∼150-250 µm), which represents the peri-injured area and Region 3 representing the “area of interest” (∼200-400 µm) in which all the experiments were performed. (B) High magnification confocal images of somatosensory cortex immunostained with a marker for astrocytes (GFAP, green) or microglia (Iba-1, red) in contra and ipsilateral S1 cortex. On the right, insets from individual microglia cells displaying the distinct phenotypes. Scale bar: 5 µm. Yellow arrow represents the “bushy” microglia phenotype represented on the right. Sections were counterstained with DAPI (blue) to illustrate nuclei. (C) GFAP^+^ cells quantification in the S1 cortex from sham (n = 6) and TBI mice treated with vehicle (n = 8) or TAK242 (n = 8; 3 mg/kg). (D) Iba1^+^ cells quantification in the S1 cortex from sham (n = 5) and TBI (n = 5 veh, n = 6 TBI TAK) mice according to the microglial inflammatory phenotype. (E) Left: schematic illustration of Sholl analysis performed to determine microglial phenotype. Right: number of intersections of the microglia branches with the designed circles and distance from soma of the branches in each group (sham + contralateral vehicle = 27 cells/11 mice; TBI veh, n = 12 cells/5 mice; TBI TAK242, n = 21 cells/5 mice). (F) Quantification of primary branches of microglial cells in the S1 cortex after TBI treatment with vehicle or TAK242 (3 mg/kg) from the same sample in E. Values are mean ± SEM. Two-way ANOVA, *Bonferroni’s* post-hoc test, * *p* < 0.05.

By quantifying microglia-released cytokines in TBI brains (Fig. 2), we also found increased TNFα (Fig. 2B) and IL-1β (Fig. 2C), which was reverted by TAK242 administration (TNFα, one-way ANOVA F_(4,29)_ = 5.25, p = 0.0026; IL-1β, one-way ANOVA F_(4,28)_ = 8.96, p < 0.0001), corroborating previous findings showing that TLR4 regulates the inflammatory response of microglia. Together, these data indicate that the inhibition of TLR4-pathway activation after our TBI model modulates both microglia activation and astrocyte reactivity.

**Figure 2.**
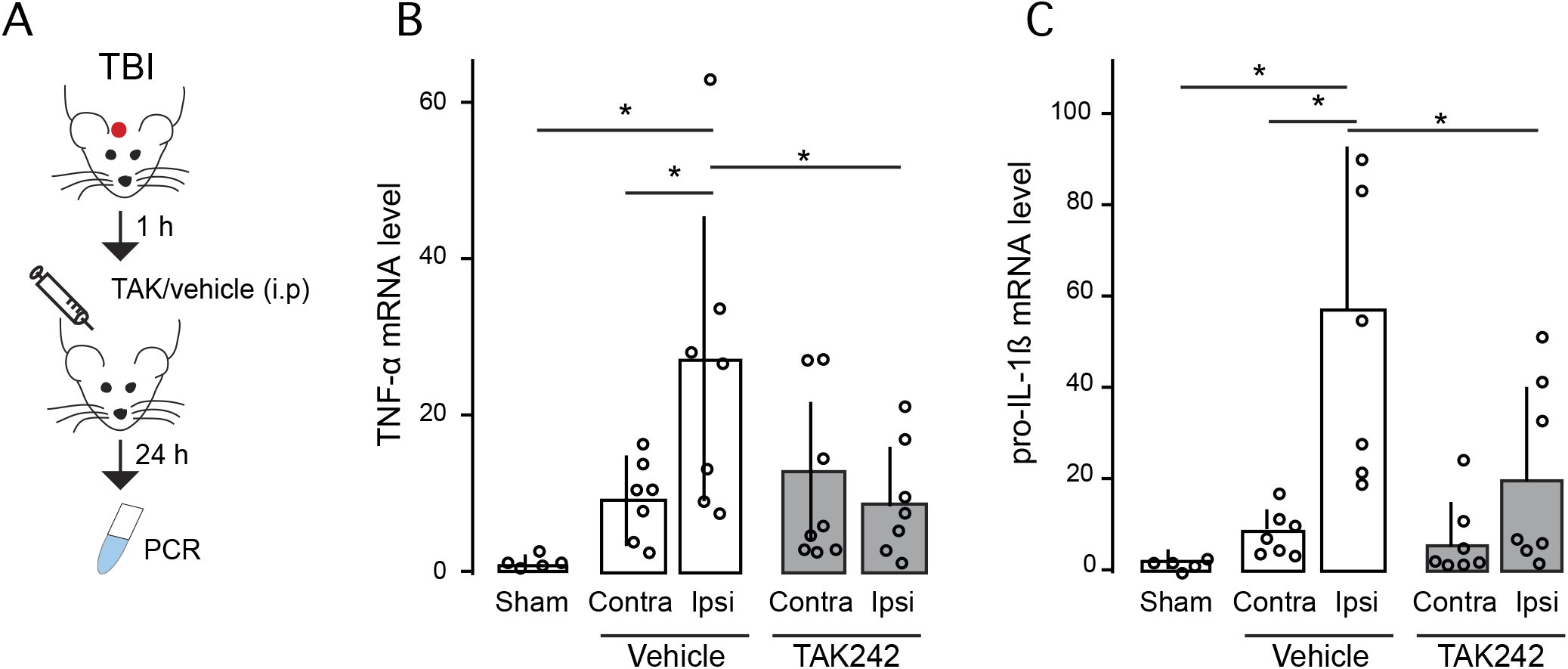
TBI-induced raise of pro-inflammatory cytokines is reverted by TAK242 treatment. (A) The schematic diagram shows the experimental strategy and the timing of the pharmacological treatment. TAK242 or vehicle were i.p. injected 1 hour after the injury. mRNA levels of (B) TNF-α (sham = 5, vehicle-treated TBI = 7, TAK-treated TBI = 8) and (C) pro-IL-1β genes (sham = 5, vehicle-treated TBI = 7, TAK-treated TBI = 7). Values are mean ± SEM. One-way ANOVA, *Bonferroni’s* post hoc test, * *p* < 0.05.

### Acute TBI recruits cortical astrocytes with somatic Ca^2+^ transients via TLR4 activation

Reactive astrocytes are known to result in a spectrum of morphological and molecular changes that influence injury progression and recovery (Nikolakopoulou et al.2016; Ohsumi et al.2010); however, little is known about the changes in functional activity of reactive astrocytes that could affect further synaptic stability and neuronal networks. Using epifluorescence microscope imaging, we examined spontaneous somatic Ca^2+^ transients from astrocytes loaded with the astrocytic marker SR101 and the intracellular calcium dye Fluo4-AM localized at 200-400 µm of the damaged area of TBI and sham mice (Fig. 3A-B). Spontaneous Ca^2+^ transients were defined as a change in fluorescence intensity relative to baseline. The proportion of “active astrocytes” showing 1 or more Ca^2+^ transients during the 1-min recording period were markedly increased in the ipsilateral TBI cortex at 1 d.a.i. (Fig. 3D, one-way ANOVA F_(4,25)_ = 9.627, p < 0.001). These effects were independent of neuronal activity, as TTX bath application did not prevent the increase in the active astrocytic population (two-way ANOVA *hemisphere* F_(1,18)_ = 40.48, p < 0.001, *treatment* F_(1,18)_ = 3.2, p = 0.09; *interaction* F_(1,18)_ = 0.007, p = 0.934; n = 5). No changes in the frequency of spontaneous Ca^2+^ transients in astrocytes were observed (Fig. 3D, one-way ANOVA F_(4,25)_ = 1.81, p = 0.157), suggesting that acute TBI leads to functional changes in cortical astrocytes observed as an increase in the number of responsive cells nearby the injury rather than an intrinsic increase in the astrocytic Ca^2+^ transients or extension of intracellular Ca^2+^ network. We next tested to which degree the recruitment of active astrocytes in the peri-injured area was dependent on TLR4-pathway (Fig. 3C). TAK242 treatment drastically decreased the number of active astrocytes in the ipsilateral cortex with no changes in the Ca^2+^ oscillations frequency (Fig. 3D). These results indicate that TLR4 activation plays a crucial role in triggering anatomical and functional changes in cortical astrocytes following TBI.

**Figure 3:**
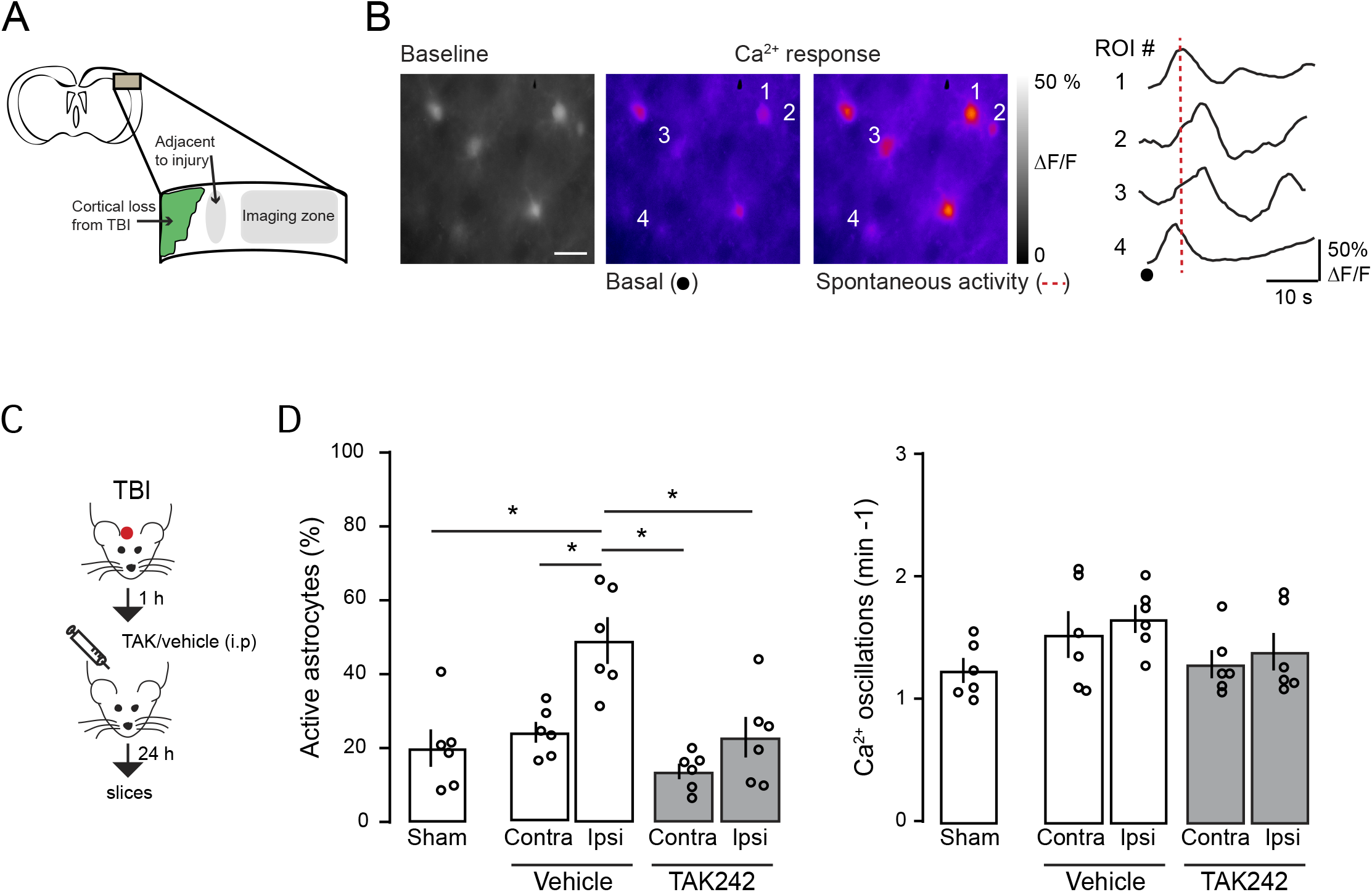
Traumatic brain injury recruits active astrocytes through TLR4 activation. (A) Schematic view of the Ca^2+^ imaging experiments showing the cortical area where astrocyte activity is registered. (B) Left: fluorescent intensities of a TBI S1 cortex loaded with Fluo4-AM (1 µg/µL) and heat maps of maximal ΔF/F before and during a spontaneous activity indicated in the rightmost figure. Color scale indicates normalized changes in fluorescence during spontaneous activity. Scale bar: 10 µm. Right: time course of the spontaneous Ca^2+^ oscillations from astrocytes indicated on the left. (C) Schematic diagram showing the experimental strategy for the injury and the pharmacological treatment. (D) Left: proportion of active astrocytes that showed at least one Ca^2+^ event during a 1-min recording period in sham (n = 6 animals/432 ROIs), vehicle (n = 6 animals, contra: 432 ROIs and ipsi: 480 ROIs) or TAK242 (n = 6, contra: 336 ROIs and ipsi: 720 ROIs) group. Right: Corresponding frequency of Ca^2+^ transients. Values are mean ± SEM. One-way ANOVA, *Bonferroni’s* post-hoc test, * *p* < 0.05.

### TLR4-induced astrocyte reactivity affects blood-brain barrier damage after TBI

Next, we investigated the consequences of the TLR4-dependent recruitment of astrocytes in the TBI pathophysiology. We first examined the BBB integrity, which is highly regulated by astrocytes and disrupted following TBI (Jhaet al.2018; Liauw et al.2008; Min et al.2015; Villapol et al.2014). S100β is an astrocytic marker released into the plasma following BBB disruption (i.e. higher the severity of BBB disruption, more S100β in serum is found). One-way ANOVA revealed a statistical significance in the amount of S100β after TBI and following TAK242 treatment (Fig. 4A, one-way ANOVA F_(2,16)_ = 17.46, p < 0.0001). A Bonferroni *post-hoc* test showed increased amount of S100β in the serum collected 1 d.a.i, indicating elevated astrocyte activity and/or damage. TAK242 treatment decreased ∼0.4-fold S100β levels, corroborating a role for TLR4 activation in the astrocyte modulation. We then injected Evans Blue dye intraperitoneally in TBI-mice and assessed BBB integrity by measuring the amount of dye leakage into the brain parenchyma (Fig. 4B). Dye leakage was mostly detected around the injury in the ipsilateral cortex, with very low leakage levels in the contralateral cortex (Fig. 4B and 4C). Blockage of TLR4-pathway decreased dye-staining as showed by the two-way ANOVA showing significant effects for hemisphere (F_(1,24)_ = 6.93, p = 0.0146), treatment (F_(1,24)_ = 76.86, p < 0.001) and interaction (F_(1,24)_ = 7.3, p = 0.012). To test whether the effect of TLR4 on BBB integrity was directly related to astrocytes, we used the inositol 1,4,5-trisphosphate receptor type 2 knockout mice (*Itpr2^−/−^*, herein referred to as IP_3_R2^−/^). IP_3_R2 is selectively expressed in astrocytes, and its genetic deletion abolishes somatic Ca^2+^ transients in these mice (Petravicz et al.2014; Srinivasan et al.2015). Two-way ANOVA showed that ipsilateral dye leakage was also found increased after TBI in IP_3_R2^−/−^ animals (Fig. 4C, hemisphere F_(1,22)_ = 88.0, p < 0.001), and it was even higher when compared to WT extravasation (unpaired *t-test* p = 0.0051), indicating that decreased astrocyte activity enhances BBB rupture following TBI. TAK242 treatment in IP_3_R2^−/−^ mice did not lessen Evans Blue extravasation (Fig. 4C, two-way ANOVA, treatment F_(1,22)_ = 0.051, p = 0.479 and hemisphere vs treatment F_(1,22)_ = 0.901, p = 0.352), and it was significantly different from values obtained in TAK-treated WT mice (avg ± sem: WT, 23.7 ± 4.55; IP_3_R2^−/−^ 53.67 ± 10.05; unpaired *t-test* p = 0.0264), indicating that TLR4-pathway partially regulates BBB rupture through modulation of astrocyte activity.

**Figure 4:**
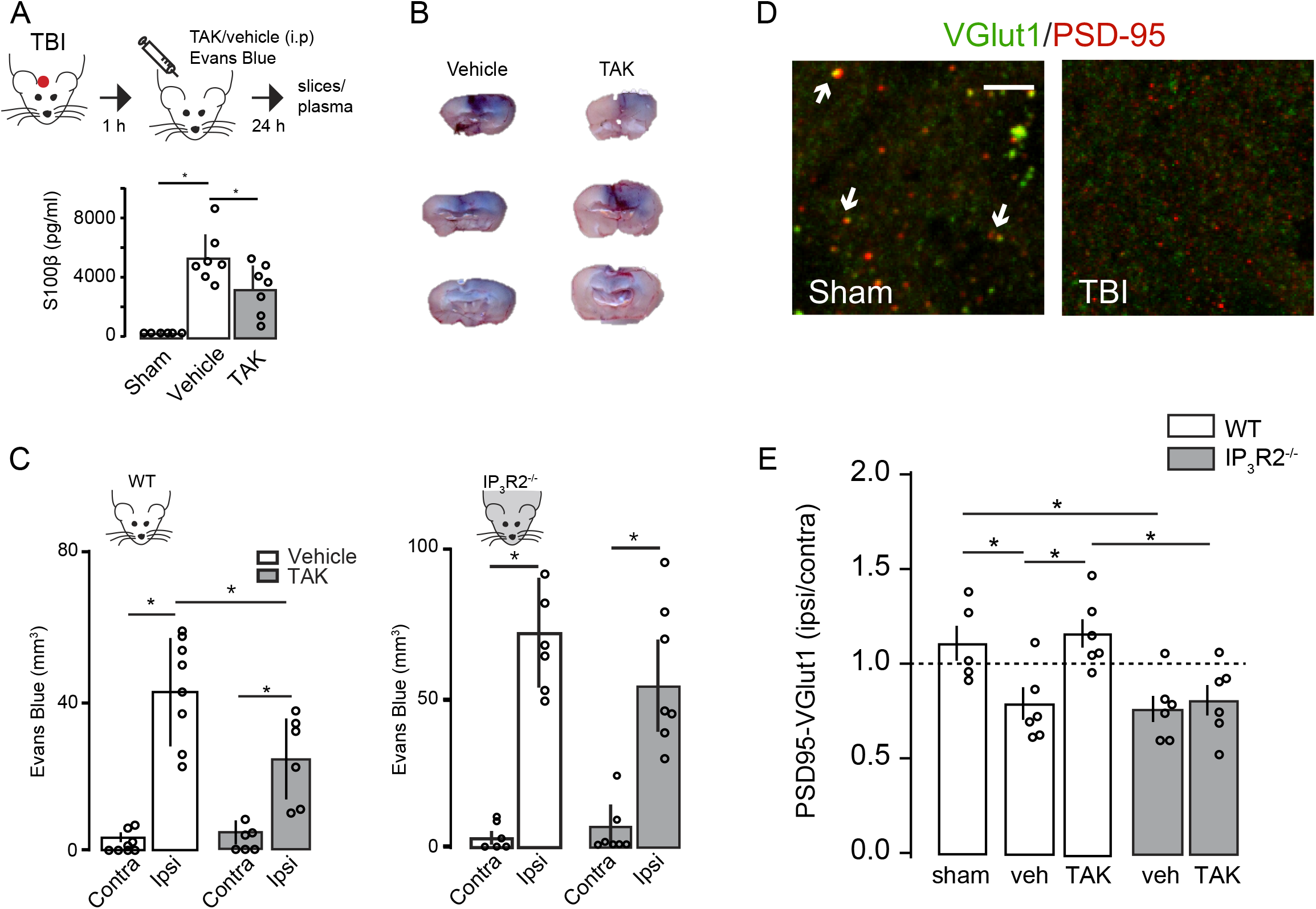
TLR4 antagonism improves synaptic loss and prevents blood-brain barrier damage after TBI. (A) Top: Schematic diagram showing the experimental strategy. Bottom: Bar graph showing the effect of vehicle or TAK242 administration on the levels of S100-β in plasma (sham = 5; TBI veh = 7; TBI TAK = 7). (B) Illustration of three distinct coronal sections through the TBI center showing the Evans Blue distribution throughout a vehicle or TAK242-treated mouse brain. (C) Blood-brain barrier integrity assessed by Evans Blue extravasations (mm^3^) 24 h after TBI in the contralateral and ipsilateral brain of WT mice (vehicle, n = 8; TAK242, n = 6) or IP_3_R2^−/−^ mice (vehicle, n = 6; TAK242, n = 7). (D) Representative confocal images of synapses (white arrowheads) indicated by colocalization of presynaptic (VGlut1, green) and postsynaptic (PSD-95, red) markers in sham and TBI ipsilateral S1 cortex. Scale bar: 5 µm. (E) Quantitative analysis showing the number of synaptic puncta 24 h after TBI in WT (sham n = 5; TBI veh n = 6; TBI TAK242 n = 6) and IP_3_R2^−/−^ (grey bars) mice (TBI veh n = 6; TBI TAK242 n = 6). Data represented as the ratio of synapses (ipsi/contra) in which values < 1 means synaptic loss in the ipsilateral cortex. Values are mean ± SEM. One-way ANOVA, Bonferroni’s post-hoc test, * *p* < 0.05, for Figure 4A and 4E. Two-way ANOVA, Bonferroni’s post-hoc test, * *p* < 0.05, for Figures 4D.

### Changes in the number of synaptic puncta density in S1 cortex after TBI

Microglia and astrocytes influence neuronal transmission through the regulation of synapses and the modulation of neuronal network (Araque et al.2014; Béchade et al.2013; Perea et al.2009; Wake et al.2009). To further investigate the pathophysiological consequences of TLR4-pathway activation, we quantified the synaptic density surrounding the injured cortex by double immunostaining for pre- and post-synaptic proteins, i.e. the number of colocalized pre-synaptic VGlut1 and post-synaptic PSD-95 puncta, at 1 d.a.i. We defined the normalized synaptic density as the synaptic ratio in the surrounding TBI area to that in the homotopic non-injured contralateral cortex to correct for any endogenous differences in the case of transgenic mice. Our results showed that ipsilateral cortex had significantly less PSD-95/VGlut1 punctas compared to sham (Fig. 4D-E, one-way ANOVA F_(4,24)_ = 5.78, p = 0.002). TAK242 treatment partially reduced the synaptic loss indicating that blockage of TLR4-pathway shortly after TBI either restores synaptic level or inhibits synaptic loss. We next quantified the number of synaptic puncta in TBI brain from IP_3_R2^−/−^ mice to test whether the neuroprotective effect on synapses observed following TAK242 treatment was exerted by TLR4-induced astrocyte activation. IP_3_R2^−/−^ mice had decreased synaptic loss following TBI, indicating that decreased astrocyte activity did not prevent synaptic loss after TBI. In addition, a partial synaptic recover was not observed after pharmacological treatment of IP_3_R2^−/−^ mice with TAK242 (Fig. 4E), suggesting that modulation of TLR4-pathway after TBI only affects synapses in the presence of active astrocytes.

### TSP-1 astrocyte release regulates synaptic number in the sub-acute phase of TBI

Since the mechanisms controlling microglia and astrocyte activation are known to have distinct phases from hours-to-days after injury (Susarla et al.2014), we determined the anatomical and functional changes of TLR4-induced astrocyte activation 7 d.a.i (Fig. 5A). Calcium imaging experiments from TBI cortex revealed a similar number of active astrocytes (one-way ANOVA F_(2,12)_ = 0.26, p = 0.775) and Ca^2+^ oscillation frequency (one-way ANOVA F_(2,12)_ = 1.1, p = 0.364) as compared to those sham or contralateral cortex (Fig. 5B). Increased number of GFAP^+^ astrocytes was still observed 7 d.a.i. (Fig. 5C, one-way ANOVA F(_2,15_) = 4.75, p = 0.025), but such increase was smaller than those found 1 d.a.i (unpaired *t-test* p = 0.049). Second, we determined the changes in microglia morphology by using both phenotype counting and Sholl analysis (Fig. 5D). No differences were observed neither in the number of non-inflammatory (one-way ANOVA F_(2,12)_ = 1.503, p = 0.26) or pro-inflammatory microglia (one-way ANOVA F_(2,12)_ = 1.21, p = 0.32) as well as Sholl analysis compared to sham 7 d.a.i. (unpaired t-test primary branches p = 1). Finally, we determined the levels of synaptic co-localization and found similar synaptic number in TBI mice to those sham (Fig. 5E), indicating a synaptic recovery in a sub-acute phase. The later corroborates previous anatomical and electrophysiological findings showing an initial synaptic loss and a decreased neuronal excitability followed by its gradual homeostatic recovery after ischemic stroke or TBI in mice (Carron et al.2016a; Ding et al.2011; Johnstone et al.2015; Johnstone et al.2013; Liauw et al.2008). Altogether, our data indicate that the recovery of microglial non-inflammatory phenotype and astrocyte activity 7 d.a.i. is accompanied by a recovery in the number of synapses.

**Figure 5:**
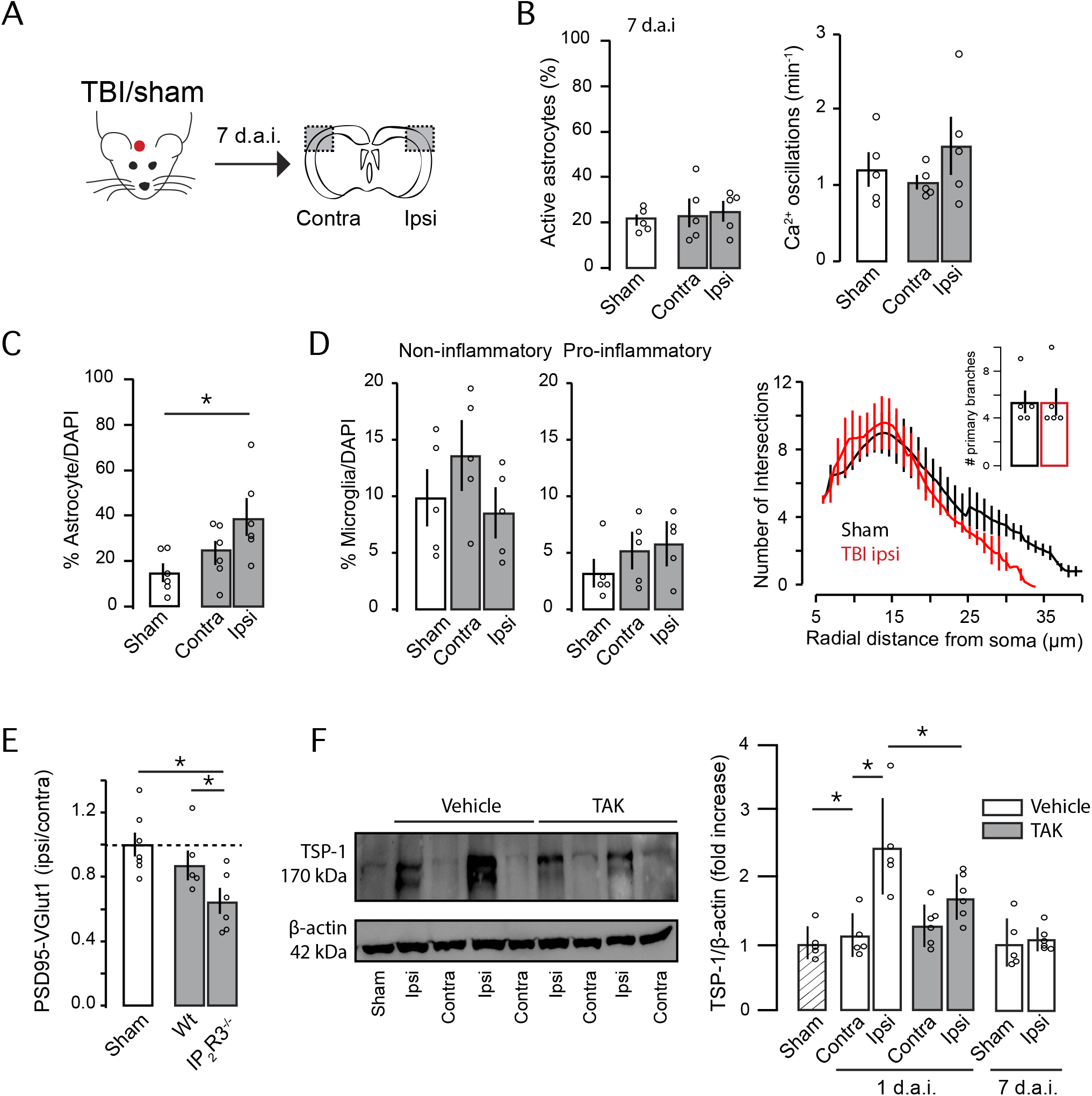
Sub-acute recovery of astrocyte activity and synaptic density in TBI brain 7 days after TBI. (A) Schematic diagram showing the experimental strategy. (B) Percentage of active astrocytes (left) and number of Ca^2+^ oscillations (right) in contra and ipsilateral cortex of sham (n = 5) or TBI mice (n = 5) 7 days after the injury. (C) Percentage of GFAP^+^ astrocytes in contra and ipsilateral cortex of TBI mice 7 days after injury (n = 6). (D) Quantitative assessment of microglial phenotypes (Iba-1 staining) in the S1 cortex from sham and TBI mice. Sholl Analysis of microglia morphology and primary branches (inset; sham = 5, TBI = 5). (E) Quantitative analysis of synaptic puncta density 7 days after TBI in WT and IP_3_R2−/− (sham, n = 7; WT, n = 5; IP_3_R2−/−, n = 6). (F) Left, original western blot of TSP-1 and actin from sham, two vehicles (Veh) and two treated mice (TAK242). Right, semi-quantitative analysis of TSP-1 immunoreactivity of TSP-1 after TBI (for 24 h mice: sham, n = 5, vehicle; n = 5, TAK242, n = 6. For 7 d.a.i. sham = 5; TBI ipsi = 6). Values are mean ± SEM. One-way ANOVA, Bonferroni’s post-hoc test, * *p* < 0.05.

We next sought to identify the potential molecular mechanism that causally links astrocyte activation to synaptic regulation at 7 d.a.i. We focused on TSP-1, which is an astrocyte-secreted soluble protein that is necessary and sufficient to promote synaptic formation during neuronal development or following brain damage (Christopherson et al.2005; Liauw et al.2008). Increased TSP-1 expression was found in ipsilateral cortex at 1 d.a.i., which returned to basal levels at 7 d.a.i. (Fig. 5F, one-way ANOVA F_(6,31)_ = 9.55, p < 0.001) (Cheng et al.2017; Liauw et al.2008). Blockage of TLR4-signaling significantly reduced TSP-1 expression to similar levels to sham or contralateral tissues, indicating that TLR4 activation controls TSP-1 release from astrocytes following TBI. To investigate whether the transient up-regulation of TSP-1 protein within the first few hours was linked to the synaptic remodelling at 7 d.a.i, we used the IP_3_R2^−/−^ mice known to have reduced release of TSP-1 after injuries (Kim et al.2016). Synaptic puncta number in IP_3_R2^−/−^ TBI brain remained decreased at 7 d.a.i. (Fig. 5E, one-way ANOVA F_(2,15)_ = 5.72, p = 0.014), indicating the important role of astrocyte activation in the homeostatic regulation of synapses in a sub-acute phase through the release of TSP-1.

### Blockage of TLR-pathway improves functional outcome after TBI

Synaptic damage is thought to be a major contributor to neurological symptoms and behavioral deficits following brain injuries. Given the significant effect on synapses and BBB integrity, we explored the possibility that TLR4-induced astrocyte activation might also affect neurological behaviour associated with TBI. We therefore investigated whether TAK242 treatment improved functional outcome by subjecting animals to a neurological severity score (NSS), a battery of behavioral tests specifically developed to assess motor and neurobehavioral outcome in TBI mice (Fig. 6, Flierl et al.2009). One-hour following TBI, mice had a NSS score around 7 points, indicating high functional deficits. Twenty-four hours later, the performance of TBI mice gradually recovered reaching values close to 5 (Fig. 6B vehicle). TAK242-treated mice at one hour after injury exhibited a better NSS score 24-hour later when compared to those animals treated only with the vehicle (two-way ANOVA effects for treatment F_(1,38)_ = 3.49, p = 0.06, time F_(1,38)_ = 56.46, p < 0.001 and interaction F_(1,38)_ = 4.11, p = 0.049). To determine whether the improvement of neurological deficits after TAK was dependent on astrocyte activation, we used IP_3_R2^−/−^ mice. Two-way ANOVA showed no effect for treatment (F_1,20_ = 0.023), time (F_1,20_ = 2.78) and interaction (F_1,20_ = 0.207), indicating that IP_3_R2^−/−^ mice did not display neurological improvement after TAK242 treatment. These data demonstrate that blockage of the TLR4-induced astrocyte activation has beneficial effects on early neurological motor tests following TBI.

**Figure 6:**
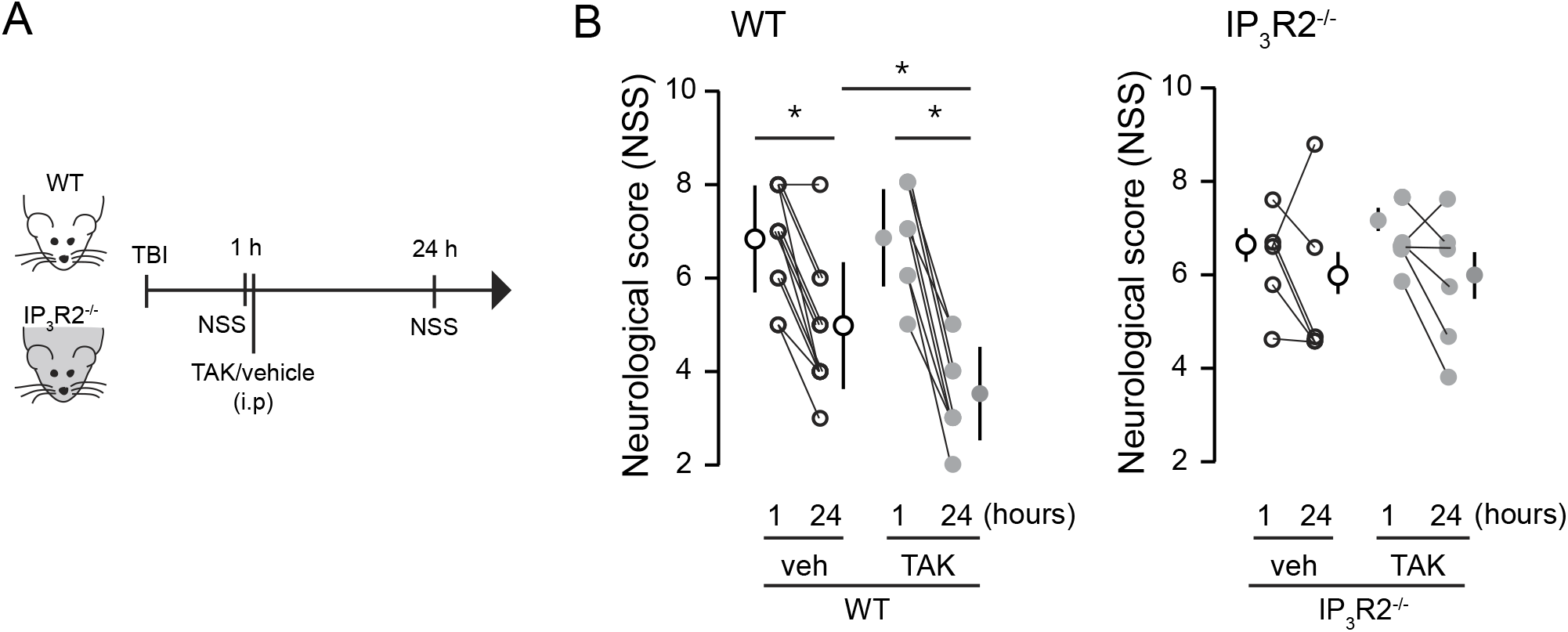
Neurological and behavioral outcome is improved by TAK242 in WT mice but not in IP_3_R2 mice. (A) Schematic diagram showing the experimental strategy. (C) Effects of TLR4 pharmacological blockade on neurological severity scores (NSS) in (left) WT (vehicle, n = 12; TAK, n = 9) and (right) IP_3_R2^−/−^ mice (vehicle, n = 6; TAK, n = 6). NSS was evaluated at 1 and 24 h after the injury. Values are mean ± SEM. Two- way ANOVA, Bonferroni’s post-hoc test, * *p* < 0.05.

In order to determine the cell target of the TAK242 pharmacological inhibition, we used immune cell characterization by flow cytometry to analyse the presence of peripheral/resident immune cells exhibiting TLR4-receptors in the ipsilateral cortex of TBI mice (Fig. 7A-C). At 1 d.a.i., TLR4 expression was mainly observed in myeloid cells, being infiltrating-monocytes and microglia the two main subpopulations with the highest percentage of TLR4^+^ signal. These data pointed to a crucial role of resident/infiltrated myeloid cells in the observed pharmacological effect of TAK242 on BBB, synapse remodelling and behavioural outcome 24 h after TBI.

## Discussion and Conclusions

We identified a fine-tune interaction which modulates the synaptic number and BBB damage with pathophysiologic relevance for long-term plasticity in TBI. We show that acute TBI recruits active astrocytes surrounding the injury in a TLR4-dependent manner (Fig. 1-3) with detrimental effects on synapses, BBB and neurological outcome (Fig. 4-6). On the other hand, TBI-induced astrocyte activation mediates the release of the astrocytic protein TSP-1 with beneficial effects on synaptic recovery a few days after injury (Fig. 5-6). Hence, pharmacological modulation of TLR4-induced astrocyte activation slow-down the damage progression and, in turn, may accelerate long-term plasticity and rebuilding of neuronal networks in the damaged area following TBI.

Microglia and infiltrating monocytes immediately respond to tissue damage by polarizing into a pro-inflammatory or into an alternative/anti-inflammatory phenotype (Davis et al.1994). Similarly, transcriptome analyses have identified two types of reactive astrocytes: A1, that up-regulate genes described to be destructive to synapses, suggesting a detrimental role, and A2, that increase neurotrophic factors and thrombospondins that promote synapse repair, suggesting a neuroprotective role (Liddelow et al.2017; Shinozaki et al.2017; Zamanian et al.2012). Hence, the inflammatory response is not an isolated response of a single cell type but is composed of a set of different resident/infiltrated cells that, as a whole, offer an orchestrated and fine-tuned response after damage. Our data evidenced that the CHI model of TBI induces a subtype of microglia and astrocyte with “harmful” effects that might correspond to the pro-inflammatory microglia and A1 astrocytes. The control of microglial inflammation exerts neuroprotective effects in several injury/disease models (Alawieh et al.2018; Donat et al.2017; Parada et al.2019) and our data indicates that the some of the acute detrimental effects of microglia after TBI, most probably through TLR4-pathway activation, is only achieved by a direct interaction with astrocytes. However, we cannot discard that some of the anti-inflammatory effects of TLR4-blocker administration might be also mediated by modulation of the inflammatory responses of infiltrated monocytes. On the other hand, the acute activation of astrocytes seems to be needed to restore synaptic levels in a sub-acute phase. Therefore, astrocyte activation has a dual role depending on the time point after injury: in an acute phase, its role is detrimental for synapses and BBB, and in the sub-acute phase is important for synaptic remodelling after TBI.

### Acute traumatic brain injury recruits active astrocytes

The induction of reactive astrocytes during the pathogenesis of brain injuries has been extensively reported by anatomical and molecular techniques (Carron et al.2016b; Kim et al.2016). In contrast, only few studies have explored the functional changes in astrocytes following CNS injuries (Kim et al.2016; Takatsuru et al.2013). By using Ca^2+^ imaging from TBI brains, we demonstrate that active cortical astrocytes are recruited in the injured area within the first few hours following TBI. No changes in the frequency of the spontaneous responses was observed, suggesting that the astrocytic network is unaffected despite the increased extracellular glutamate after TBI (Cantu et al.2015). The recruitment of active astrocytes was abolished by blocking TLR4-pathway activation after the first hour post injury, which led to an improvement in the synaptic number as well as BBB integrity. These data support the idea that TLR4-induced astrocytes activation play a detrimental role in the neuronal and vascular functions, and therefore, blockage of TLR4-pathway may be a potential therapeutic target for TBI.

### Activation of TLR4-pathway regulates synaptic puncta following TBI

Microglia and astrocytes are known to control synapses and to modulate neuronal networks after brain lesions or diseases. For example, in *in vivo* models, microglia activation of the complement cascade controls neuronal synaptic pruning (Tremblay et al.2010; Wake et al.2009); while microglia-derived molecules regulate synaptic strength and neuronal activity (Béchade et al.2013) by inducing synaptic facilitation (Clark et al.2015) and increasing the frequency of spontaneous excitatory synaptic currents (Pascual et al.2012). Independently of the mechanism used by microglia to control synaptic transmission, it is clear that microglia plays a crucial role in the control of neuronal networks. On the other hand, astrocytes also regulate synaptic transmission and neuronal homeostasis through several mechanisms (Araque et al.2014; Navarrete et al.2019; Rosa et al.2015). *In vivo* models show that astrocytic release of TSP-1/2 is associated with synapse formation and axonal outgrowth (Kim et al.2016; Liauw et al.2008); while enhanced astrocytic production and release of D-serine or ephrins are detrimental to synapses (Nikolakopoulou et al.2016; Perez et al.2017). Here, we give new evidence that astrocyte regulates neuronal synapses through the activation of TLR4-pathway in inflammatory cells following TBI. TLR4-receptors are mainly implicated in the recognition of infectious pathogens or endogenous ligands released upon injury and are the main pathway to activate microglia after TBI. However, TLR4 is also expressed in other cell types such as endothelial and peripheral immune cells (monocytes, neutrophils, granulocytes and lymphocytes) (for more see https://www.brainrnaseq.org/) that access the brain parenchyma following a robust BBB rupture (Priego & Valiente, 2019). Given that infiltrated-monocytes and microglia, but not neutrophils and lymphocytes, were the most prominent myeloid cell types expressing TLR4 under our experimental model, we cannot discard that some of the neuroprotective effects observed after systemic administration of TLR4 blocker might be also mediated by modulation of the inflammatory responses via monocytes. On the other hand, infiltrating-monocytes influence the proliferation of juxtavascular astrocytes located in the proximity of damaged area following traumatic injury (Bardehle et al., 2013; Frik et al., 2018), which could also affect synaptic remodelling through the modulation of astrocyte and microglia activation. Supporting our hypothesis, we found that synaptic lost is partially reverted by modulating TLR4-pathway activation in an astrocyte-dependent manner, since IP_3_R2^−/−^ mice fail to recover synapses after TLR4-receptors blockage. In addition, astrocytic-protein TSP-1 released following TBI seems to regulate the synaptic recovery in the sub-acute phase of the injury. Given all these evidences, our results support that control of astrocyte activation through inhibition of TLR4-pathway in inflammatory cells plays an important role in the regulation of synapses following TBI.

The effect of pharmacological blockage of TLR4 observed in this study could also be attributed to a direct effect on astrocytes since several studies have shown TLR4 expression in cultured astrocytes. However, these data is currently contradictory with some reports showing that TLR4 is expressed in both microglia and astrocytes (Du et al., Front Mol Neurosc, 2017, Lisa et al Glia 2013,), while others showing that TLR4 response in astrocytes is only mediated by previous microglia activation (Holm et al., Glia 2012). Nonetheless, several recent reports using cutting-edge genetic techniques have shown a predominant expression of TLR4 in microglia, but not in astrocytes (Zhang et al., Neuron 2016, Zhang et al., J Neurosc 2014, Cahoy et al., J Neurosc 2008). Our data using IP_3_R2^−/−^ mice support the idea that the effect observed in this study is mediated by TLR4-pathway activation either in microglia or infiltrating peripheral cells.

### Sub-acute effects of TLR4-induced astrocyte activation on synapses

Reactive astrocytes exert neuroprotective effects after TBI by limiting the extension of the injured core and by forming a glial scar (Shinozaki et al. 2017). Our data extend the beneficial effects of astrocytes to the synapse recovery in the TBI sub-acute phase. By using anatomical quantification of PSD-95/VGlut1 puncta, we demonstrate a significant recovery of the initial synaptic loss a week following injury. This physiological mechanism seems to depend on the astrocyte activation as IP_3_R2^−/−^ mice continue to present reduced synaptic number 7 d.a.i. Our data also indicates that such synaptic regulation depends on TSP-1, an astrocyte-released protein that serves to control neuronal synapses after brain damage (Christopherson et al.2005). In our experiments, TSP-1 is transiently upregulated 1 d.a.i, but its effect on synapse is observed only a few days after injury. This data corroborates other studies suggesting that TSP-1 effects may take a several day period to be observed (Christopherson et al.2005), as it acts on the expression of proteins that participate in the formation of new synaptic proteins rather than regulating cell adhesion proteins to lock synapses into place (i.e. stabilizing role) (Liauw et al.2008). So, while neuronal cortical synapses are rapidly lost (probably due to direct neuronal death), TLR4-induced astrocyte activation increases TSP-1 release in the first few hours to trigger the expression of neuronal synaptic proteins later on, leading to a partial synaptic recovery 7 d.a.i. Interestingly, the synaptic recovery can be speed-up by an order of days after pharmacological TLR4-blockage. Hence, TLR4-inhibition of astrocyte activation works to slow-down the progression of synaptic damage, thereby accelerating the recovery/repair of the surrounded damaged area.

### Inhibition of astrocyte activation through TLR4 blockage improves BBB after TBI

Both microglia and astrocytes physically interact with brain endothelial blood-vessels influencing BBB integrity (Burda et al.2016). Here, we found that while TLR4-blockage attenuates BBB rupture after TBI, this effect is not observed in IP_3_R2^−/−^ mice, indicating a direct effect of astrocyte and a strong relationship between inflammatory cells and astrocyte (Donat et al.2017; Jha et al.2018). The astrocytic detrimental effects on BBB after TBI could be explained by the release of vasoactive endothelial growth factor (Argaw et al.2009) or apolipoprotein E (Bell et al.2012) shown to exacerbate BBB damage. However, astrocytes are also known to release molecules that act beneficially on endothelia to reduce BBB permeability and promote its repair after CNS injury including Sonic hedgehog (Alvarez et al.2011) and retinoic acid (Mizee et al.2014). Such results divergence demonstrates that astrocytes are pivotal regulators of endothelial BBB properties that act to open, maintain or restore barrier functions via specific molecular mechanisms that remain to be investigated.

## Conclusion

Our data provide new evidence showing a role of TLR4-mediated astrocyte activation in the pathophysiology of brain injuries (Fig. 8). We postulate that after a TBI, extracellular molecules that are rapidly released upon tissue damage activate TLR4-signaling to recruit active astrocytes in the peri-injured area that in turn will affect neuronal excitability by modulating synaptic number in a time-dependent manner. Shortly after the injury, astrocyte decreases synapses probably to slow-down the neuronal network and to avoid further neurotoxic damage. At the same time, active astrocytes release TSP-1 to promote synaptic recovery later on. After that, a homeostatic control of both microglia and infiltrated-monocyte activity may take place followed by a decrease in reactive/active astrocytes. Moreover, this cell-to-cell interaction also exacerbates immediate BBB breakdown probably due to the release of astrocytic proteins and pro-inflammatory cytokines known to have detrimental effects on BBB. Overall, our data show that decreasing TLR4 activation and the pro-inflammatory microglia phenotype and infiltrated-monocytes during the first few hours following a TBI could be used as a target for therapeutic treatments that may promote synaptic plasticity and improve cognitive function via astrocytic modulation.

**Figure 8:**
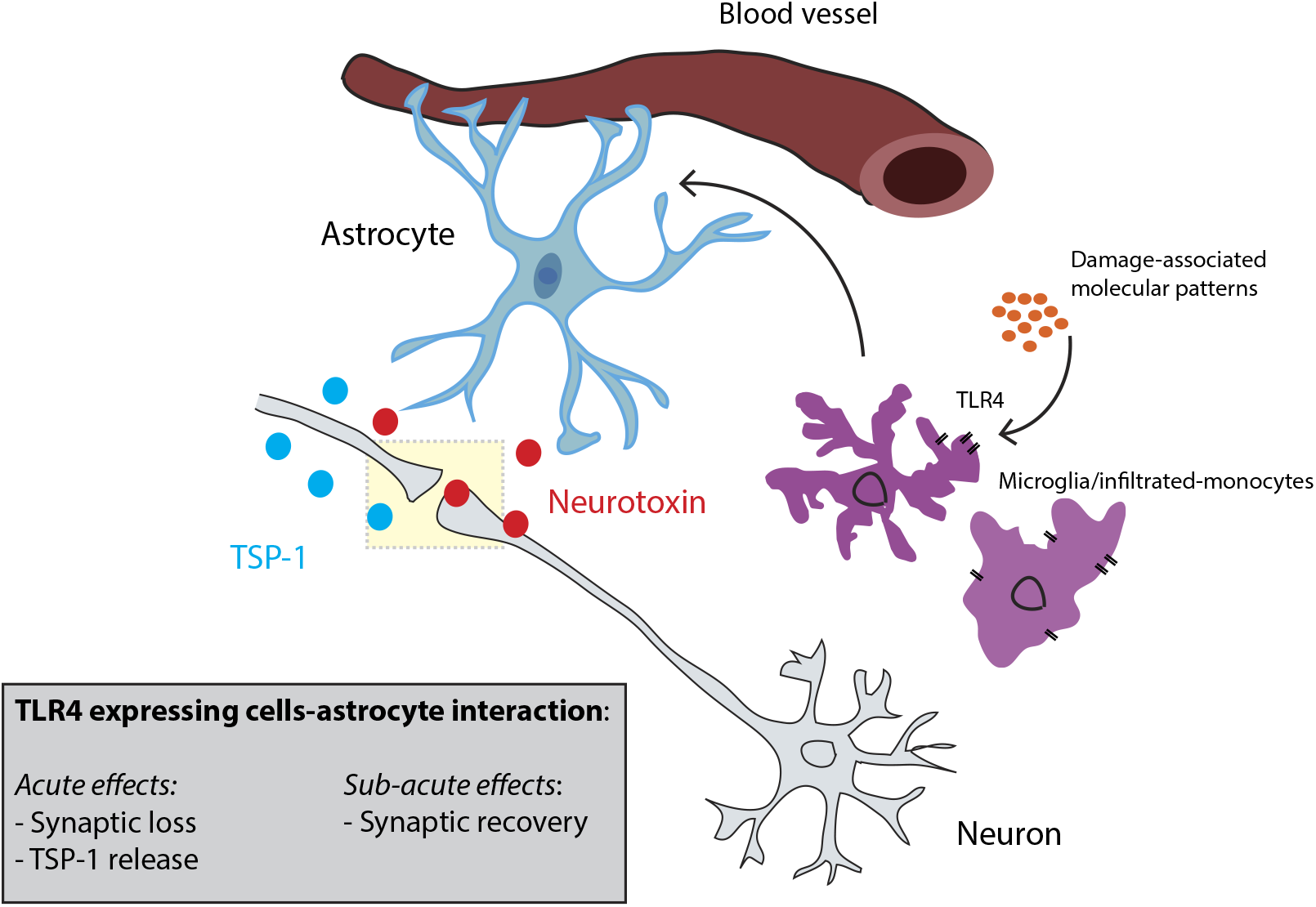
Schematic diagram of the microglia/infiltrated monocyte-astrocyte interaction following TBI. Control of astrocyte activation represents a key physiological mechanism to regulate both neuronal vasculature and synaptic number following TBI in a time-dependent manner. Shortly after the injury, astrocyte activity decreases the number of synapses probably to slow down the neuronal network and avoid further neurotoxic damage. At the same time, active astrocytes release TSP-1 to promote synaptic recovery later on. After this, a homeostatic control of microglia and infiltrated monocytes activity may take place followed by a decrease in active astrocytes and neuronal excitability. Moreover, inflammatory cell types-astrocyte interaction also exacerbates immediate BBB breakdown probably due to the release of astrocytic proteins and proinflammatory cytokines known to have detrimental effects on BBB.

## Acknowledgments

This work was supported by grants from the Instituto de Salud Carlos III (ISCIII) (Programa Miguel Servet II Grants CPII19/00005; PI16/00735; PI19/00082 to JE and PI18/00357 to DC, partially funded by FEDER-European Union “*Una manera de hacer Europa*”) and Fundación Mutua Madrileña to JE. European Union’s Horizon 2020 research and innovation programme under the Marie Sklodowska-Curie grant agreement No 794926 and Stop Fuga de Cerebros Roche Pharma to JMR. RTI2018-094887-B-I00 and RYC-2016-20414 to MN. DC,MCO and VVS are hired by SESCAM. We also thank Dr. Ju Chen (University of California, CA, San Diego, USA) and Dr. Gertrudis Perea (Cajal Institute Madrid, Madrid, Spain) for kindly providing IP3R2^−/−^ animals and Dr. Juan Aguilar (HNP, Toledo, Spain) for technical and scientific support.

## Author contributions

JMR and JE conceived and designed the study. JMR and VFA performed and analysed immunofluorescence experiments. VF performed and analysed Evans Blue extravasation experiments. JMR performed, analysed and wrote the scripts for Ca^2+^ imaging experiments. MCO, DC and VVS designed, performed, analysed and wrote the text about flow cytometry experiments. MN assisted with astrocyte imaging experiments and advised on calcium imaging analysis. APA, EFL and PNF assisted with IP_3_R2^−/−^ experiments. VF and CD performed and analysed PCR experiments. All authors revised the manuscript. JMR and JE wrote the manuscript.

